# Phosphoproteomics Reveals a Distinct Mode of Mec1/ATR Signaling in Response to DNA End Hyper-Resection

**DOI:** 10.1101/2020.04.17.028118

**Authors:** Ethan J. Sanford, Vitor M. Faça, Stephanie C. Vega, William J. Comstock, Marcus B. Smolka

## Abstract

The Mec1/ATR kinase is crucial for genome maintenance in response to a range of genotoxic insults, although how it promotes context-dependent signaling and DNA repair remains elusive. Here we uncovered a specialized mode of Mec1/ATR signaling triggered by the extensive nucleolytic processing (resection) of DNA ends. Cells lacking *RAD9*, a checkpoint activator and an inhibitor of resection, exhibit a selective increase in Mec1-dependent phosphorylation of proteins associated with single strand DNA transactions, including the ssDNA binding protein Rfa2, the translocase/ubiquitin ligase Uls1 and the HR-regulatory Sgs1-Top3-Rmi1 (STR) complex. Extensive Mec1-dependent phosphorylation of the STR complex, mostly on the Sgs1 helicase subunit, promotes an interaction between STR and the DNA repair scaffolding protein Dpb11. Fusion of Sgs1 to phosphopeptide-binding domains of Dpb11 strongly impairs HR-mediated repair, supporting a model whereby Mec1 signaling regulates STR upon hyper-resection to influence recombination outcomes. Overall, the identification of a distinct mode of Mec1 signaling triggered by hyper-resection highlights the multi-faceted action of this kinase in the coordination of checkpoint signaling and HR-mediated DNA repair.

## INTRODUCTION

In eukaryotes, PI3K-like kinases (PIKKs) play essential roles in the sensing of DNA damage and the coordination of cell cycle checkpoint activation and DNA repair mechanisms to maintain genome stability (Savitsky et al., 1995; Shiloh, 2003). In *Saccharomyces cerevisiae*, the Mec1 PIKK (human ATR) plays a central role in coordinating the response to DNA lesions that result in single stranded DNA (ssDNA) exposure, including stalled replication forks and recessed double strand breaks (DSBs) (Cha & Kleckner, 2002; Deshpande et al., 2017; Segurado & Diffley, 2008; Tercero, Longhese, & Diffley, 2003). Mec1 senses ssDNA exposure through the recognition of replication protein A (RPA)-bound ssDNA via its obligate cofactor Ddc2 (human ATRIP) (Deshpande et al., 2017; Paciotti, Clerici, Lucchini, & Longhese, 2000; Zou & Elledge, 2003). Once recruited to ssDNA, activation of Mec1 requires the action of Mec1 activating proteins that contain Mec1 activation domains (MADs). Three Mec1 activators have been identified in budding yeast: Ddc1, a member of the 9-1-1 PCNA-like clamp; Dna2, a flap endonuclease with established roles in DNA end resection, and Dpb11, a multi-BRCT domain containing scaffolding protein (Kumar & Burgers, 2013; Majka, Niedziela-Majka, & Burgers, 2006; Mordes, Nam, & Cortez, 2008; Navadgi-Patil & Burgers, 2009; Wanrooij & Burgers, 2015). The substrate specificity of the activators is distinct—Dna2 recognizes DNA flaps whereas Ddc1-Dpb11 is loaded onto 5’ dsDNA-ssDNA junctions produced directly, but not exclusively, as a result of DNA end resection (Majka, Binz, Wold, & Burgers, 2006; Stewart, Campbell, & Bambara, 2009).

In its canonical mode of action, Mec1 activates the downstream kinase Rad53 to mediate a DNA damage checkpoint response that arrests the cell cycle and reshapes the transcriptional and replication programs (Desany, Alcasabas, Bachant, & Elledge, 1998; Huang, Zhou, & Elledge, 1998; Lanz, Dibitetto, & Smolka, 2019; Seeber, Dion, & Gasser, 2013). Mec1 also phosphorylates a range of other targets to mediate checkpoint-independent responses (BastosdeOliveira et al., 2015; Lanz et al., 2018). Cells lacking *MEC1*, but not cells lacking *RAD53*, have extraordinarily high rates of gross chromosomal rearrangements (Myung, Datta, & Kolodner, 2001), pointing to a crucial checkpoint-independent role for Mec1 in genome maintenance. Despite the importance, the mechanisms by which Mec1 suppresses genomic instabilities remain incompletely understood.

Recent biochemical and genetic evidence points to key roles for Mec1/ATR in the control of homologous recombination (HR) (Barlow & Rothstein, 2009; Dion, Kalck, Horigome, Towbin, & Gasser, 2012; Flott et al., 2011; Ullal, Vilella-Mitjana, Jarmuz, & Aragón, 2011; W.-L. Toh et al., 2010), a multi-step DNA repair process essential for maintaining genome integrity during the S and G2 phases of the cell cycle (Moynahan & Jasin, 2010). The essential first step of HR is resection, the 5’-3’ nucleolytic degradation of DNA ends. Resection is followed by strand invasion, DNA synthesis, end ligation, and the processing of recombination intermediates such as Holliday junctions (Mimitou & Symington, 2009; West et al., 2016). Given its complexity, HR requires exquisite regulation to prevent deleterious outcomes. For example, strand invasion can occur at the wrong locus, leading to non-reciprocal translocations or other complex chromosomal rearrangements (Putnam & Kolodner, 2017).

Mec1 has been reported to control distinct steps in HR. Most notably, Mec1 plays both inhibitory and activating roles in the control of DNA end resection. Mec1 phosphorylation of histone H2A (γH2AX in mammals) and of the Ddc1 subunit of the 9-1-1 complex assembles a ternary complex involving the resection antagonist Rad9 at DNA lesions (Fig. 1A). Mec1-dependent stabilization of Rad9 at DNA lesions antagonizes resection (Clerici, Trovesi, Galbiati, Lucchini, & Longhese, 2014; Liu et al., 2017). Mec1 also phosphorylates the DNA repair scaffold Slx4, which counteracts Rad9 recruitment and therefore alleviates the block in resection (Dibitetto et al., 2015; Lanz et al., 2019; Liu et al., 2017; Ohouo, Bastos De Oliveira, Liu, Ma, & Smolka, 2013). Therefore, Mec1 balances anti- and pro-resection outcomes for proper resection control. In addition, Mec1 phosphorylates the recombinase protein Rad51 to control HR through inactivation of Rad51 ATP hydrolysis and DNA binding activities (Flott et al., 2011). It remains unknown whether Mec1 controls additional steps or proteins for HR control.

**Figure 1.**
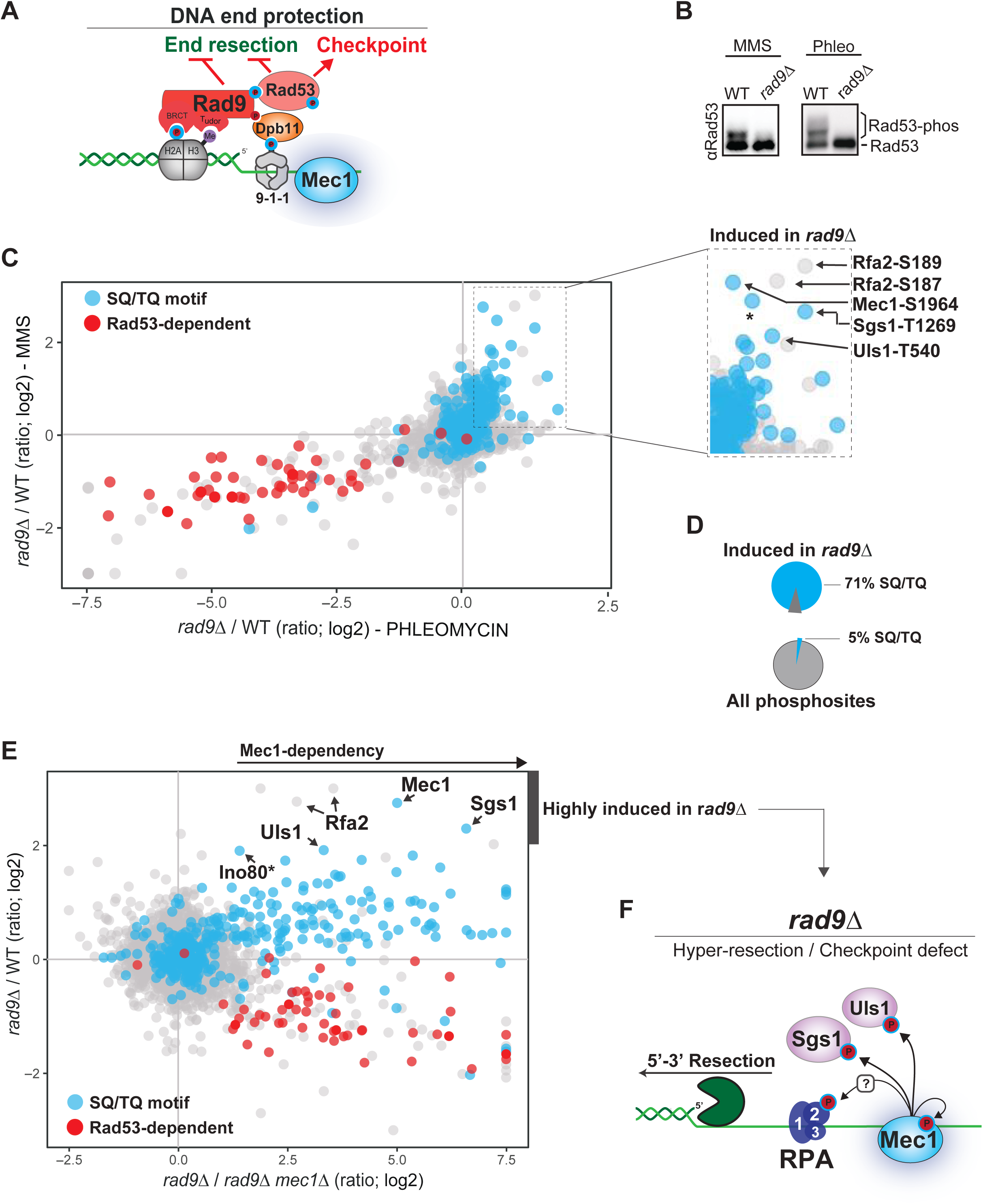
A Distinct Mode of Mec1/ATR Signaling in *rad9Δ* cells. A. Model showing the role of Rad9 in activating the Rad53-dependent checkpoint and preventing DNA end resection. B. *rad9Δ* cells fail to activate the Rad53 checkpoint in the presence of either the DNA alkylating drug MMS (0.02%) or the radiomimetic drug phleomycin (40ug/mL). C. Quantitative phosphoproteomics dataset showing that S/T-Q phosphorylation is enriched in cells lacking RAD9. Among the most highly enriched sites are Sgs1 T1269, Uls1 T540, Mec1 S1964, and 2 S-N sites in Rfa2. This response is similar in both the presence of 40ug/mL phleomycin (x axis) and 0.02% MMS (y axis). Note downregulation of Rad53 signaling (red dots) due to absence of RAD9. D. Pie chart showing that S/T-Q phosphorylation comprises a large fraction of *rad9Δ*-induced phosphorylation (71%), though only accounting for a small fraction (5%) of the entire dataset. E. Quantitative phosphoproteomics dataset showing that most phosphorylation events induced in *rad9Δ* cells are dependent on Mec1. F. Model depicting Mec1-dependent phosphorylation of Sgs1, Uls1, and Rfa2 in response to hyper-resection in *rad9Δ* cells.

Dpb11, in addition to its role as an activator of Mec1 signaling, also functions as a key scaffolding protein to mediate the formation of ternary complexes involved in the DNA damage response. Dpb11 contains four BRCA1 C-terminus-like (BRCT) domains that act, often pairwise, to bind phosphorylated amino acid residues on client proteins (Cussiol, Jablonowski, Yimit, Brown, & Smolka, 2015; Wardlaw, Carr, & Oliver, 2014).

To date, a number of Dpb11 interacting partners with roles in the DNA damage response have been identified, including the checkpoint mediator Rad9, the DNA repair scaffold Slx4, and the nuclease Mus81-Mms4 (Gritenaite et al., 2014; Ohouo, Bastos de Oliveira, Almeida, & Smolka, 2010; Pfander & Diffley, 2011). By activating Mec1 and assembling these complexes, Dpb11 dictates the spatiotemporal dynamics of Mec1 signaling and its outcomes. Early in the response, Dpb11 bound to the checkpoint adaptor protein Rad9 mediates the transduction of signaling from Mec1 to the downstream checkpoint kinase Rad53, thereby establishing a cell cycle checkpoint response (Fig. 1A) (Pfander & Diffley, 2011; Schwartz et al., 2002). Dpb11-mediated stabilization of Rad9 at 5’ recessed ends of DNA lesions is also important to promote Rad9’s function in blocking DNA end resection (Lazzaro et al., 2008; Liu et al., 2017; Villa, Bonetti, Carraro, & Longhese, 2018). In the absence of Rad9, cells fail to properly activate the DNA-damage checkpoint and DNA ends undergo rapid end resection (Lazzaro et al., 2008). Cells lacking *RAD9* exhibit a greater incidence of non-allelic recombination, and this defect is consistent with the fact that loss of *RAD9* frequently produces synergistic increases in chromosomal rearrangement in mutants lacking factors responsible for the regulation of homology-directed repair (Fasullo, Bennett, Ahching, & Koudelik, 1998; Nielsen, Bentsen, Andersen, Gasser, & Bjergbaek, 2013).

To explore additional roles for Mec1 in HR control, we monitored DNA damage signaling in cells lacking Rad9, which lack proper checkpoint signaling and DNA end protection, and therefore undergo extensive DNA end resection. Using phosphoproteomics, we find that *rad9Δ* cells exposed to DNA damage exhibit a specialized mode of Mec1 signaling converging toward the phosphorylation of proteins involved in ssDNA-associated transactions, including the ssDNA binding protein RPA, the Sgs1 helicase and the Uls1 translocase. In *rad9Δ* cells, Mec1 mediates an interaction between the STR (Sgs1-Top3-Rmi1) complex and the Dpb11 scaffold. We propose that, upon hyper-resection, Mec1 signaling regulates STR to influence recombination outcomes. Overall, the identification of a distinct mode of Mec1 signaling triggered by hyper-resection highlights the multi-faceted action of this kinase in the coordination of checkpoint signaling and HR-mediated DNA repair.

## RESULTS

### Loss of Rad9 stimulates a specialized mode of Mec1 signaling

To explore checkpoint-independent roles of Mec1 downstream of resection control, we monitored Mec1-dependent signaling events in wild-type and *rad9Δ* cells treated with the DNA alkylating agent methylmethanesulfonate (MMS) or the radiomimetic drug phleomycin. Rad9 is known to play a key role in activating the downstream kinase Rad53 upon formation of ssDNA gaps by MMS or DSBs by phleomycin treatment (Figs. 1A and B). Quantitative phosphoproteomics confirmed that Rad53 phosphorylation as well as Rad53-dependent phosphorylation events are impaired in *rad9*Δ cells (Fig. 1C and Supplemental Table S1). Strikingly, the results show a set of phosphorylation events induced in *rad9*Δ cells, mostly on the preferential SQ/TQ motif for Mec1 or Tel1 phosphorylation (Figs. 1C-D and Supplemental Table S1). Analysis comparing *rad9Δ* cells with or without Mec1 confirmed that most of the SQ/TQ phosphorylation stimulated in *rad9Δ* cells is indeed dependent on Mec1 (Fig. 1E and Supplemental Table S1). Lack of *RAD9* stimulated Mec1 autophosphorylation at S1964, but not autophosphorylation at S38, suggesting that in *rad9*Δ cells Mec1 adopted a distinct mode of activation/signaling compared to its mode of signaling in wild-type cells. The proteins with the strongest increases in Mec1-dependent phosphorylation in *rad9Δ* cells have been implicated in ssDNA-associated DNA repair transactions, including the ssDNA binding protein Rfa2, the Sgs1 helicase and the Uls1 translocase/ubiquitin ligase (Figs. 1C, 1E-F and Supplemental Table S1). Phosphorylation on Rfa2 was not on the SQ/TQ consensus, but on SN and SA sites, raising the possibility that Mec1 is either able to phosphorylate these other consensus motifs in the context of *rad9*Δ or that such sites are phosphorylated indirectly through the control of another kinase. Overall, our unbiased quantitative phosphoproteomic analyses reveal that lack of *RAD9* triggers a specialized mode of Mec1 signaling targeting proteins associated with HR-mediated DNA repair. Since Rad9 is important to protect DNA ends by preventing resection, and because the targets are hyper-phosphorylated in *rad9*Δ cells are involved in ssDNA-associated repair transactions, the results suggest that this may represent a mode of Mec1 signaling triggered by the over-exposure of ssDNA.

### The 9-1-1 complex plays a more prominent role than Dna2 in Mec1 activation in *rad9Δ* cells

Mec1 can be activated by two independent pathways—either by the 9-1-1 complex (formed by Ddc1-Rad17-Mec3 in yeast) in conjunction with the Dpb11 scaffold, or by Dna2 (Kumar & Burgers, 2013; Mordes et al., 2008; Navadgi-Patil & Burgers, 2009; Wanrooij & Burgers, 2015). We sought to determine the contribution of each of these modes of Mec1 activation to the Mec1 signaling response stimulated in *rad9Δ* cells. We used quantitative MS to determine the effect of deleting *DDC1*, which impairs 9-1-1/Dpb11-mediated Mec1 activation, or introducing the *DNA2*^*WYAA*^ mutation, which fails to activate *DNA2*-dependent Mec1 signaling (Kumar & Burgers, 2013). Both mutants were analyzed in the context of cells lacking *RAD9*. Whereas the *DNA2*^*WYAA*^ mutation had minimal effect on most Mec1-dependent phosphorylation (Fig. 2A, C, and Supplemental Table S2), deletion of *DDC1* resulted in stronger impairment of most Mec1-dependent signaling induced by MMS in *rad9Δ* cells. Similarly, phosphorylation of Sgs1 by Mec1 was not altered by the *DNA2*^*WYAA*^ mutation but was reduced by lack of *DDC1* (Fig. 2C and Supplemental Table S2). The reduction of detected SGS1 phosphorylation sites was not drastic, suggesting that Dna2-mediated Mec1 activation or another mode of Mec1 signaling may partially compensate for loss of DDC1. Still, the reduction in *ddc1Δ* cells was comparable in magnitude to the increase in Sgs1 phosphorylation observed in *rad9Δ* cells, suggesting that the stimulation of the phosphorylation observed in *rad9Δ* cells is principally induced upon DNA damage via the 9-1-1-dependent pathway of Mec1 activation.

**Figure 2.**
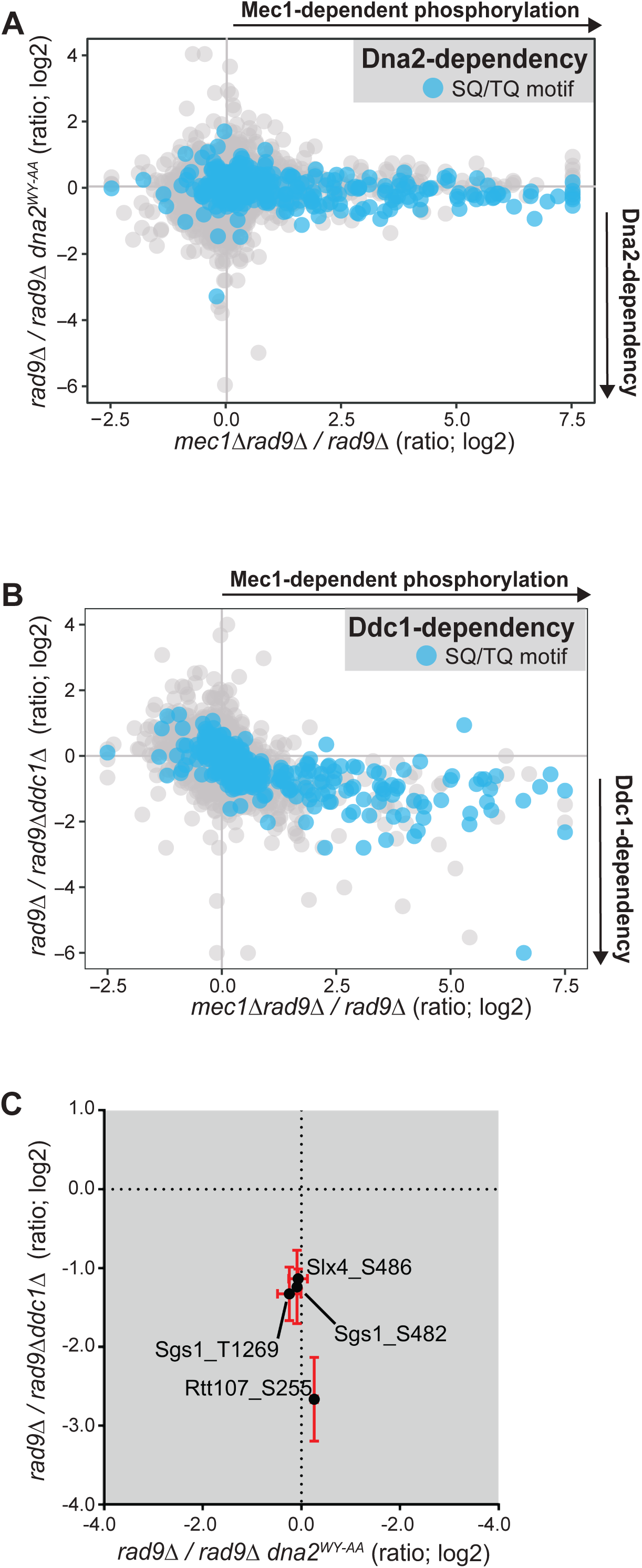
The 9-1-1 Subunit Ddc1 Plays a More Prominent Role in Activation of Mec1 in Response to *rad9Δ* than Dna2. A. Quantitative phosphoproteomic analysis defining the role of Dna2 (WY-AA mutation impairs Mec1-activating function of Dna2) in the set of Mec1-dependent phosphorylation in *rad9Δ* cells. Dna2 dependency (y axis) is plotted against Mec1 dependency (x axis). B. Quantitative phosphoproteomic analysis defining the role of Ddc1 in the set of Mec1-dependent phosphorylation in *rad9Δ* cells. Ddc1 dependency (y axis) is plotted against Mec1 dependency (x axis). C. Plot comparing Dna2 and Ddc1 dependency of selected Mec1 substrates.

### The STR complex undergoes extensive Mec1-dependent phosphorylation in *rad9Δ* cells

Sgs1 is a well-established regulator of homology-directed repair (Ashton, Mankouri, Heidenblut, McHugh, & Hickson, 2011; Bermúdez-López et al., 2016; Campos-Doerfler, Syed, & Schmidt, 2018; Chiolo et al., 2005; Klein & Symington, 2012; Lo et al., 2006; Mirzaei, Syed, Kennedy, & Schmidt, 2011). Given the key roles for Sgs1 in multiple steps of HR control, including resection, heteroduplex rejection and joint molecule dissolution, we decided to further investigate Mec1-dependent phosphorylation of Sgs1. Data from a recently deposited phosphoproteome database reveal that Sgs1 is phosphorylated at multiple sites (Fig. 3A) (bioRxiv DOI: 10.1101/700070). In addition to the sites detected in our phosphoproteomic experiments (Fig. 1), we noticed that most of the previously identified phosphorylation sites in Sgs1 cluster at the protein’s N-terminus, which contains a large unstructured region previously reported to engage in a range of protein-protein interactions (Bjergbaek, Cobb, Tsai-Pflugfelder, & Gasser, 2005; Chiolo et al., 2005; Hegnauer et al., 2012). Analysis of a previously reported Sgs1 truncation spanning the first 647 N-terminal amino acids (Sgs1^1-647^; (Mirzaei et al., 2011) revealed that upon MMS treatment, the Sgs1 N-terminus undergoes a stronger MMS-induced gel mobility shift in *rad9Δ* cells than in wild-type cells (Fig. 3B, left). Consistent with the finding that Sgs1 undergoes Mec1-dependent hyper-phosphorylation in *rad9Δ* cells (Fig. 1), the gel mobility shift of the Sgs1^1-647^ truncation in *rad9Δ* cells was severely reduced in cells lacking Mec1 (Fig. 3B, right). These results confirm that Sgs1 becomes preferentially phosphorylated in a Mec1-dependent manner in cells lacking *RAD9* and reveal that in addition to phosphorylation of threonine 1269 identified in our phosphoproteomic analysis, the N-terminus of Sgs1 also undergoes extensive of Mec1-dependent phosphoregulation.

**Figure 3.**
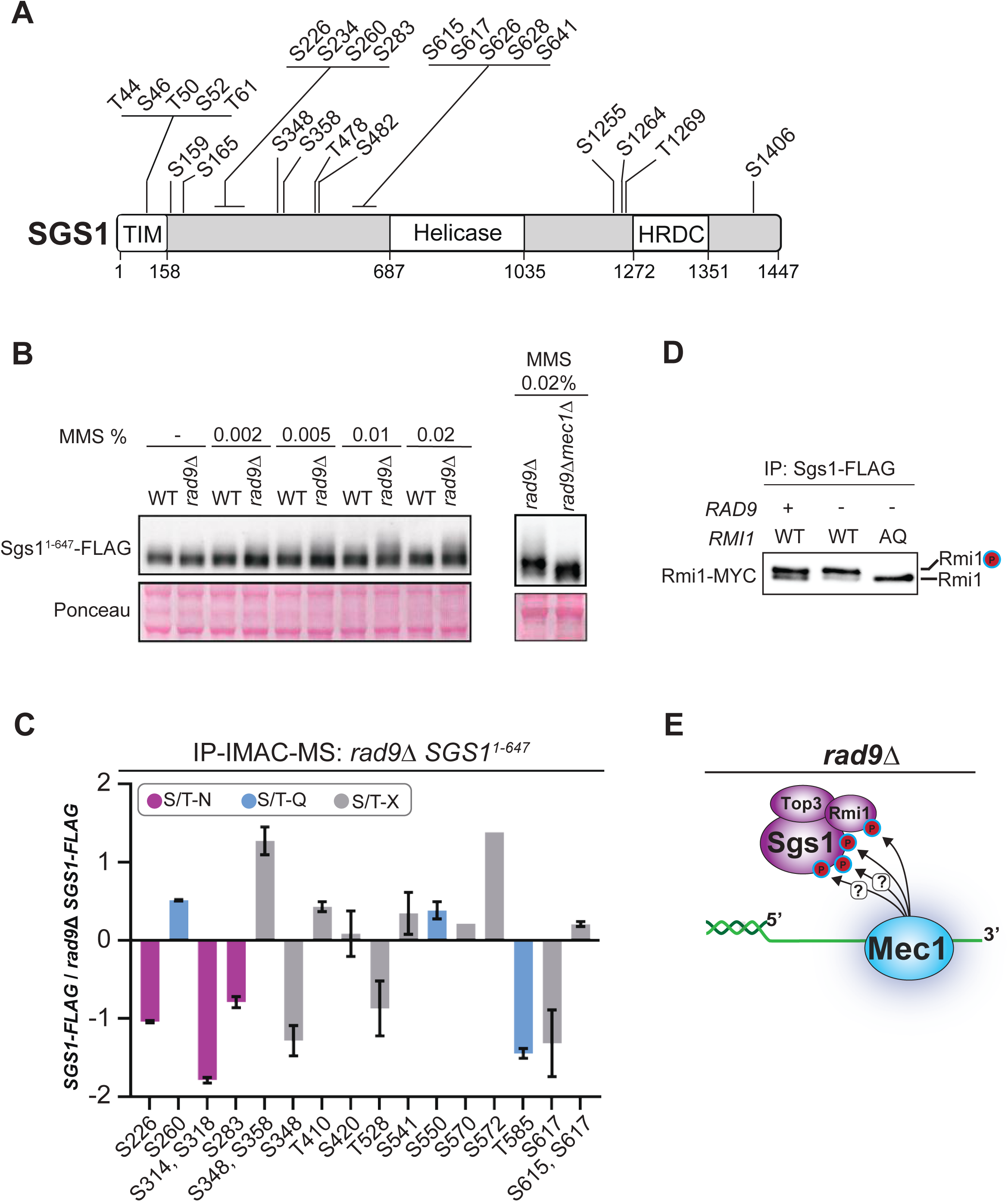
The STR Complex is Phosphorylated at Multiple Sites in Response to Hyper-Resection. A. Schematics representing data of all Sgs1 phosphorylation sites detected in a large scale, high coverage, phosphoproteomic dataset. B. Immunoblot analysis of Sgs1 N-terminus (amino acids 1-647) from cells treated with increasing doses of the DNA alkylating drug MMS in the absence of RAD9 and/or MEC1. C. Quantitative mass spectrometry analysis of phosphopeptides from affinity purified Sgs1 N-terminal fragment expressed in *rad9Δ* or wild-type MMS-treated cells. Error bars represent standard deviation. D. Rmi1, a component of the Sgs1-Top3-Rmi1 (STR) complex, is hyper-phosphorylated on its 3 S/T-Q motifs in response to *rad9Δ*. E. Model for Mec1-dependent phosphorylation of the STR complex.

Next, we performed quantitative MS analysis of the N-terminal region of Sgs1 (residues 1-647) in order to map *rad9*Δ-dependent N-terminal phosphorylation using quantitative mass spectrometry. Our analysis revealed that, while not all detected S/T-Q phosphorylation events were induced in cells lacking *RAD9*, threonine 585 phosphorylation (a TQ motif) was induced in the absence of *RAD9* (Fig. 3C and Supplemental Table S2). In addition, multiple phosphorylation sites in the S-N motif within a 100 amino acid window on the N-terminus of Sgs1 were highly induced in *rad9*Δ cells (Fig. 3C and Supplemental Table S2), as well as phosphorylation on residues 348, 528 and 617, which are not in in S/T-Q or S/T-N motifs. Therefore, our analyses indicate that the pattern of Sgs1 phosphorylation induced in *rad9*Δ cells is complex and includes non-S/T-Q motifs. Moreover, analysis of the primary sequence of the STR complex component RMI1 revealed that this small protein contains three S/T-Q motifs. To determine if Rmi1 was hyperphosphorylated in *rad9*Δ cells in the same manner as Sgs1, we analyzed by CoIP the fraction of Rmi1 bound to Sgs1 (Fig. 3D). While Rmi1 detectability was obscured in the input fraction due to a contaminating band at approximately the same molecular weight as the tagged protein, Rmi1 appears as a doublet when immunoprecipitated with Sgs1. We interpret the lower molecular weight band to be non-phosphorylated Rmi1, and the higher molecular weight band to be a phosphorylated species of Rmi1. Interestingly, the stoichiometry of Rmi1 phosphorylation changes in response to *rad9*Δ such that the intensity of the lower molecular weight band is reduced, indicating an increase in the phosphorylated species of the protein. Mutation of the three S/T-Q motifs to alanine (AQ mutant) abolishes the gel shift, consistent with these sites being targeted by Mec1. Overall, these results reveal an intricate pattern of Mec1-dependent multi-phosphorylation of the STR complex in cells lacking Rad9, which could involve additional downstream kinases (Fig. 3E).

### Sgs1 binds to the Dpb11 scaffold in cells lacking *RAD9*

Since the phosphorylation of Sgs1 by Mec1 was dependent on Ddc1-mediated Mec1 activation, we reasoned that Sgs1 might be brought in proximity to Mec1 through binding to Dpb11, a scaffolding protein that interacts with Ddc1 and previously reported to recruit known Mec1 targets, such as Slx4 and Rad9, to sites of DNA damage (Ohouo et al., 2010; Pfander & Diffley, 2011). Therefore, we affinity purified Dpb11 overexpressed in wild-type or *rad9*Δ cell lines and used mass spectrometry to identify interacting partners (Figs. 4 A-C and Supplemental Table S3). Expectedly, known Dpb11 interacting partners such as Ddc1, Sld2, Rtt107, and Slx4 copurified with Dpb11 regardless of the status of *RAD9*. By contrast, Sgs1, Top3, and Rmi1, components of the STR complex, copurified with Dpb11 only when *RAD9* was deleted (Figs. 4 C-D). To narrow down the region of Sgs1 that was responsible for binding to Dpb11, we tested whether the N-terminus of Sgs1 (amino acids 1-647; Fig. 4E), previously shown to mediate various protein-protein interactions (Bjergbaek et al., 2005; Chiolo et al., 2005; Hegnauer et al., 2012), could bind to Dpb11. As with the full-length protein, the N-terminal 647 residues of Sgs1 pulled down Dpb11 in cells lacking the checkpoint adaptor *RAD9*, but the interaction was much reduced when *RAD9* was present (Fig. 4F). Notably, the N-terminal 647 amino acid fragment of Sgs1 fused to a 3x FLAG tag was more stable than full length Sgs1, and more amenable to our coimmunoprecipitation experiments. Taken together, these findings uncover a previously undescribed interaction between Sgs1 and the Dpb11 scaffold and suggest a model whereby, in *rad9Δ* cells, Dpb11 bound to Ddc1 recruits the STR complex in proximity to Mec1 (Fig. 4G).

**Figure 4.**
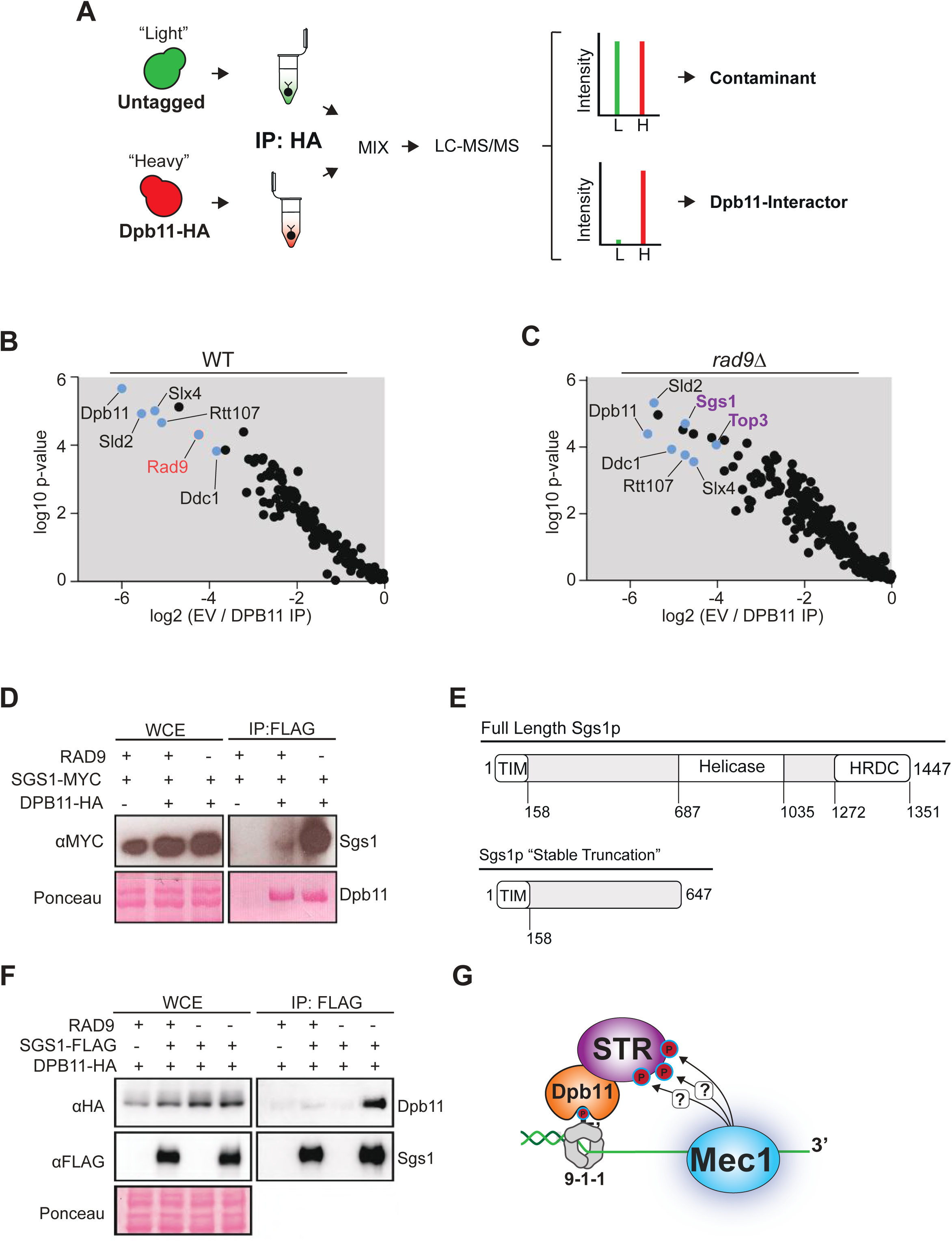
Dpb11 Interacts with Sgs1 in *rad9Δ* cells. A. Workflow of the SILAC-based quantitative mass spectrometry method for identifying Dpb11 interactions. B. Identification of Dpb11 interacting proteins in wild-type cells using workflow shown in (A). Dpb11 was overexpressed using the *ADH1* promoter. P value calculated using Mann-Whitney u-test. C. Identification of Dpb11 interacting proteins in *rad9Δ* cells using workflow shown in a. Dpb11 was overexpressed using the *ADH1* promoter. P value calculated using Mann-Whitney u-test. D. Immunoblot showing co-immunoprecipitation between Dpb11 overexpressed from an ADH1 promoter and Sgs1 expressed from its endogenous locus. E. Schematic of the Sgs1 N-terminal truncation used in this study. F. Immunoblot showing co-immunoprecipitation of Dpb11 expressed at its endogenous locus and the Sgs1 N-terminus tag-truncated with a 5xFLAG tag and expressed at its endogenous locus. G. Hypothetical depiction of the Dpb11-STR complex engaged at 5’ recessed ends. See text for details.

### Requirements for assembly of the Dpb11-STR Complex

Next, we investigated the architecture of the Dpb11-Sgs1 interaction using a combination of affinity-purification quantitative mass spectrometry and conventional co-immunoprecipitation. We used the Rad9-Dpb11 and Slx4-Dpb11 interaction architecture as a model. In these complexes, Dpb11 binds to p-Thr 602 on the Ddc1 subunit of the 9-1-1 clamp via Dpb11 BRCT domains 3 and 4. Ddc1 T602 phosphorylation is Mec1-dependent (Puddu et al., 2008), and upon engagement of the C-terminal BRCT domains of Dpb11 with the 9-1-1 clamp, the pair of N-terminal BRCTs (1 and 2) is free to engage phosphorylated interacting partners such as Rad9 and Slx4 (Cussiol et al., 2015; Pfander & Diffley, 2011).

Like the Rad9-Dpb11 and Slx4-Dpb11 complexes, the Sgs1-Dpb11 interaction was at least partially dependent on the N-terminal pair of BRCT domains in Dpb11, as evidenced by the reduction in interaction observed when the Dpb11-K55A mutant (which partially impairs BRCT 1, and binding via BRCT1/2; (Cussiol et al., 2015) was used in the co-affinity purifications followed by MS analysis (Fig. 5A and Supplemental Table S4). This result was corroborated by analysis of the Dpb11^K55A^ mutant by co-immunoprecipitation and western blot (Fig. 5B). Expectedly, our mass spectrometry experiment also confirmed that the Slx4-Dpb11 interaction was dependent on functional N-terminal BRCT domains of Dpb11 (Cussiol et al., 2015). In addition, the Sld2-Dpb11 interaction, which is important for DNA replication initiation (Kamimura, Masumoto, Sugino, & Araki, 1998) and has been previously reported to depend on the C-terminal BRCT domains of Dpb11(Tak, Tanaka, Endo, Kamimura, & Araki, 2006), was not disrupted in binding to the Dpb11^K55A^ protein (Fig. 5A). The interaction of Dpb11 with the Mus81-Mms4 complex was also not disrupted in the K55A mutant. Of importance, formation of the Sgs1-Dpb11 complex was also dependent on the 9-1-1 subunit Ddc1 and, specifically, on its phosphorylation by Mec1 at threonine 602 (Figs. 5C-D and Supplemental Table S4). Congruent with the requirement for Mec1, deletion of *MEC1* also impaired the Sgs1-Dpb11 interaction, while deletion of *TEL1* had no effect (Figs. 5E-F and Supplemental Table S4). These findings, together with our previous knowledge of how Dpb11 engages with the 9-1-1 complex, support a model in which Dpb11 engaged with the 9-1-1 clamp subunit Ddc1, via recognition of phosphorylated threonine 602 by BRCT3/4, interacts with the STR complex at least partially via BRCT1/2 of Dpb11 (Fig. 6A). Since both Dpb11 and Ddc1 are established activators of Mec1, the initial engagement of STR with Dpb11 may further promote the extensive multi-site pattern of STR phosphorylation by Mec1. Strikingly, the architecture of the complex and its requirement for assembly are mostly similar to what was previously reported for the Slx4-Dpb11 interaction (Cussiol et al., 2015; Ohouo et al., 2010).

**Figure 5.**
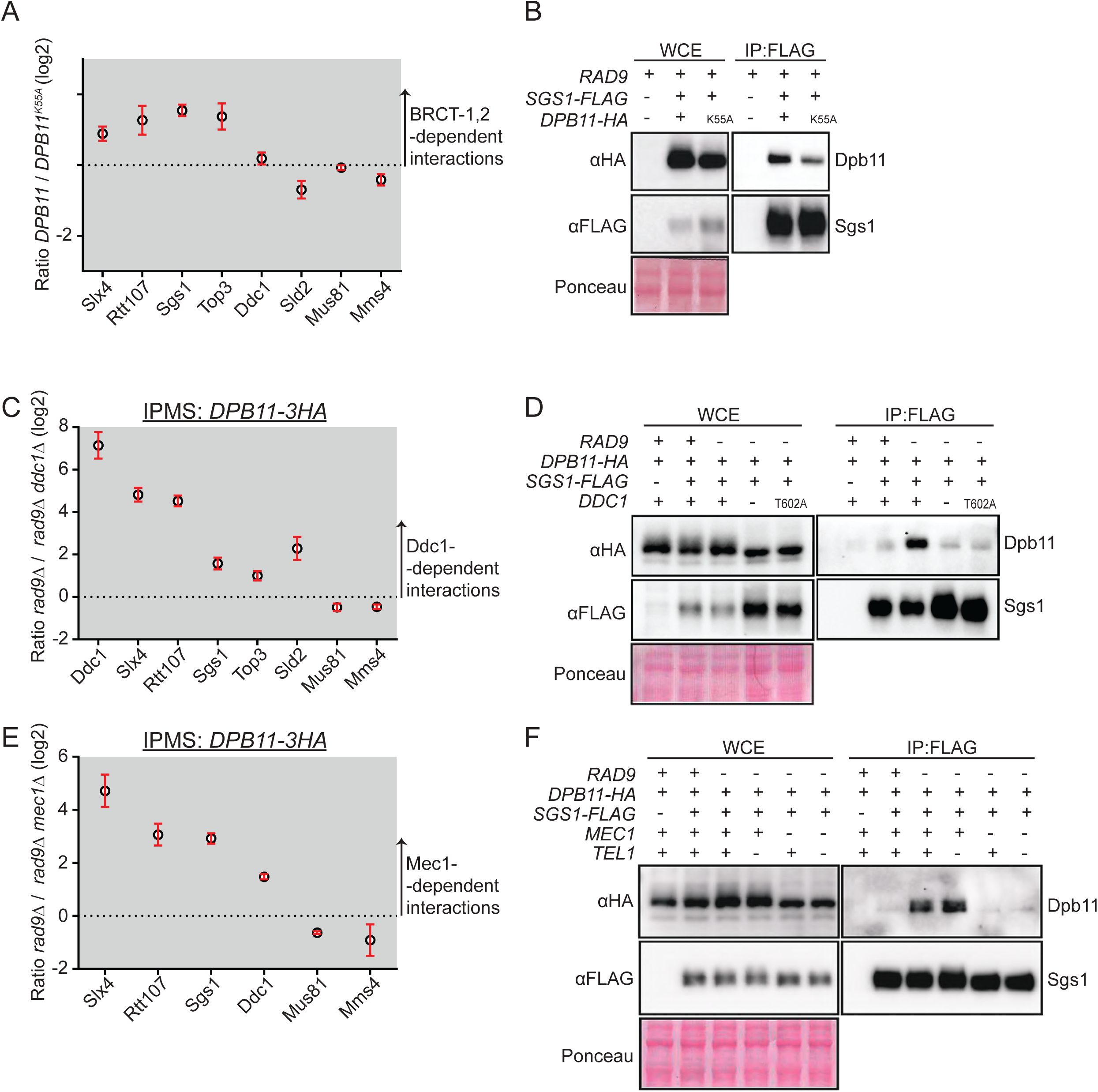
Requirements for Assembly of the Dpb11-STR Complex. A. Quantitative mass spectrometry analysis of affinity purified Dpb11 complexes, comparing wild-type Dpb11 to the *dpb11-K55A* mutant that partially impairs binding through the pair of BRCT domains 1 and 2. Cells were grown in the presence of 0.04% MMS. Error bars represent SEM. B. Immunoblots showing co-immunoprecipitation between Dpb11 and Sgs1 in the presence of 0.04% MMS with either wild-type or the *DPB11*^*K55A*^ mutant. *DPB11-3HA* was ectopically expressed from its endogenous promoter and co-immunoprecipitated with *SGS1*^*1-647-*^*FLAG* expressed from its endogenous promoter. C. Quantitative mass spectrometry analysis of affinity purified Dpb11 complexes, comparing Dpb11 purified from *rad9Δ* cells with or without the 9-1-1 subunit Ddc1. Cells were grown in the presence of 0.04% MMS. Error bars represent SEM. D. Immunoblots showing co-immunoprecipitation between Dpb11 expressed from its endogenous locus and Sgs1 tag-truncated with 5xFLAG and expressed from its endogenous locus in the presence of 0.04% MMS in either wildtype, *ddc1Δ*, or *ddc1-T602A* cell lines. E. Quantitative mass spectrometry analysis of affinity purified Dpb11 complexes, comparing Dpb11 purified from *rad9Δ* cells with our without Mec1. Cells were grown in the presence of 0.04% MMS. Error bars represent SEM. F. Immunoblots showing co-immunoprecipitation between Dpb11 and Sgs1 in the presence of 0.04% MMS in either wildtype, *mec1Δ, tel1Δ* or *mec1Δ tel1Δ* cell lines. *DPB11-3HA* was tagged at its endogenous locus and co-immunoprecipitated with *SGS1*^*1-647*^*-FLAG* tagged and truncated at its endogenous locus.

**Figure 6.**
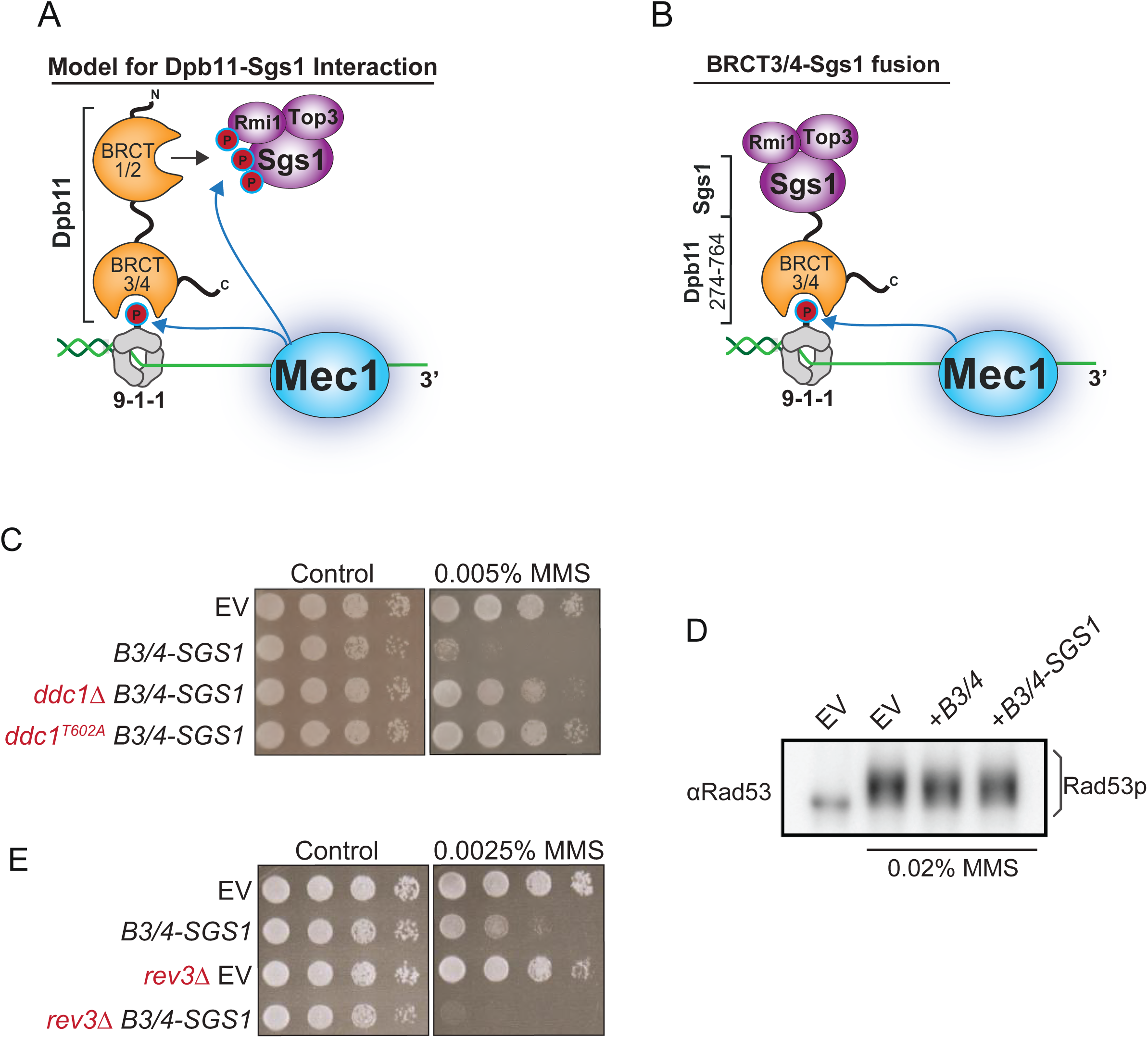
A BRCT-Sgs1 Fusion Protein Promotes MMS Sensitivity in a Ddc1-dependent Manner. A. Schematics depicting the rationale for designing a BRCT-Sgs1 chimera. The BRCT3/4-Sgs1 fusion protein is expected to engage at DNA lesions via the 9-1-1 clamp. B. Dilution assay of wild-type, *ddc1Δ*, or *ddc1-T602A* cells expressing BRCT3/4-SGS1 in the presence of 0.005% MMS. C. Immunoblot of Rad53 mobility shift assay in cells treated with 0.02% MMS expressing either BRCT3/4 of DPB11 alone or the BRCT3/4-Sgs1 chimera. D. Dilution assay of wild-type or *rev3Δ* cells expressing BRCT3/4-Sgs1 in the presence of 0.0025% MMS.

### Generation of a BRCT-Sgs1 chimeric protein for functional interrogation of the Dpb11-Sgs1 interaction

As shown in Figure 6A, we hypothesized that Dpb11 is important to recruit or stabilize, the STR complex to DNA lesions containing a loaded 9-1-1 clamp to access intermediate DNA structures formed upon the overexposure of ssDNA. To test this model and gain insights into the role of the Dpb11-Sgs1 interaction, we fused full length Sgs1 to BRCT domains 3 and 4 (BRCT3/4) of Dpb11. As shown in Figure 6A, right panel, we reasoned that this chimeric protein should promote the enhanced recruitment of STR to the 9-1-1 clamp, bypassing the need for Mec1 to promote assembly of an Sgs1-Dpb11 complex, while still relying on phosphorylation of threonine 602 in Ddc1. Expression of B3/4-Sgs1 resulted in strong sensitivity to MMS in WT cells (Fig. 6B), indicating that this chimeric protein exerts a dominant effect even in cells expressing *RAD9*. The MMS sensitivity was rescued by deletion of *DDC1* or by mutation of Ddc1 residue T602 to alanine, consistent with the model that the 9-1-1 clamp stabilizes STR at DNA lesions, which in the case of the chimeric protein, may hyper-stabilize STR leading to aberrant control of the DNA damage response. Since Sgs1 has been reported to promote Rad53 activation in response to replication stress (Hegnauer et al., 2012), we monitored whether the B3/4-Sgs1 chimera impacted DNA damage checkpoint signaling. We did not observe a change in Rad53 activation in response to MMS treatment in cells expressing the chimeric protein (Fig. 6C), suggesting that a B3/4-Sgs1 fusion protein impacts the DNA damage response independently of Rad53 activation. The STR complex plays major roles in the control of homologous recombination, including the surveillance of recombination intermediates and rejection of heteroduplexes to prevent non-allelic recombination (Ashton, Mankouri, Heidenblut, McHugh, & Hickson, 2011; Bermúdez-López et al., 2016; Campos-Doerfler, Syed, & Schmidt, 2018; Chiolo et al., 2005; Klein & Symington, 2012; Lo et al., 2006; Mirzaei, Syed, Kennedy, & Schmidt, 2011). Indeed, a major DNA repair pathway mediating MMS resistance is homologous recombination (Game & Mortimer, 1974). A plausible explanation for the MMS sensitivity observed upon B3/4-Sgs1 expression could be that it is hyper-stabilizing the STR complex at DNA lesions, which is then rejecting recombination intermediates and not allowing execution of HR-mediated DNA repair of MMS-induced lesions. Consistent with this model, expression of B3/4-Sgs1 resulted in exquisite MMS sensitivity in cells lacking REV3, a translesion synthesis factor (Fig. 6D), which represents a pathway parallel to HR for the repair of MMS-induced DNA lesions (Doles et al., 2010; Jansen, Tsaalbi-Shtylik, & de Wind, 2015). These results support the model that Dpb11 stabilizes the STR complex at DNA lesions via the 9-1-1 clamp for engagement of recombination intermediates.

### The B3/4-Sgs1 Chimera Impairs Homology-Directed Repair

To determine if the B3/4-Sgs1 fusion influences recombination outcomes we used two well-established recombination assays for monitoring break-induced replication (BIR) or gene conversion (GC) (Fig. 7) (Anand et al., 2014; Ira, Malkova, Liberi, Foiani, & Haber, 2003). Strikingly, expression of B3/4-Sgs1 severely impaired break-induced replication (Fig. 7B). The observed BIR defect was partially rescued by introducing mutations that either impair helicase activity (*hd*, helicase-defective mutation, *SGS1*^*K706A*^) or impair the ability of Sgs1 to bind to Top3 (*tim*, top3 interacting motif, *SGS1*^*Δ1-158*^) in Sgs1 (Fig. 7B). Double *hd*/*tim* mutations fully rescued by BIR defect cause by the chimera, consistent with the model that the defect is caused dominantly by STR activity (Fig. 7B). Fusing any of the 3 components of the STR complex to Dpb11 BRCT 3/4 impaired BIR to a similar extent (Fig. 7C). Fusing other helicases such as Rrm3 or Pif1 to Dpb11 BRCT 3/4 did not significantly impair BIR (Fig. 7D). Notably, fusing *H. sapiens* BLM to Dpb11 BRCT 3/4 did significantly impair BIR (Fig. 7D). Similar results were observed using the GC assay (Figs. 7E-F). Finally, we asked if the B3/4-Dpb11 fusion protein increases the stringency in single-strand annealing DNA repair using a previously published SSA reporter assay (Fig. 7G) (Sugawara, Goldfarb, Studamire, Alani, & Haber, 2004). An increased SSA stringency would be manifested in this assay as a higher ratio of AA repair efficiency to FA repair efficiency since the FA strain contains 3% sequence non-homology and is therefore increasingly subjected to Sgs1-mediated heteroduplex rejection. Indeed, expression of the B3/4-Sgs1 fusion protein led to an increase in the ratio of AA to FA repair product (Fig. 7H), consistent with the chimeric protein increasing heteroduplex rejection. Overall, the results support a model in which Dpb11 and the 9-1-1 clamp play an important role in controlling the action of STR. Since Mec1 mediates the interactions between Dpb11 with 9-1-1 and STR, our findings point to a role for Mec1 signaling in controlling recombination through the regulation of STR stabilization at DNA lesions.

**Figure 7.**
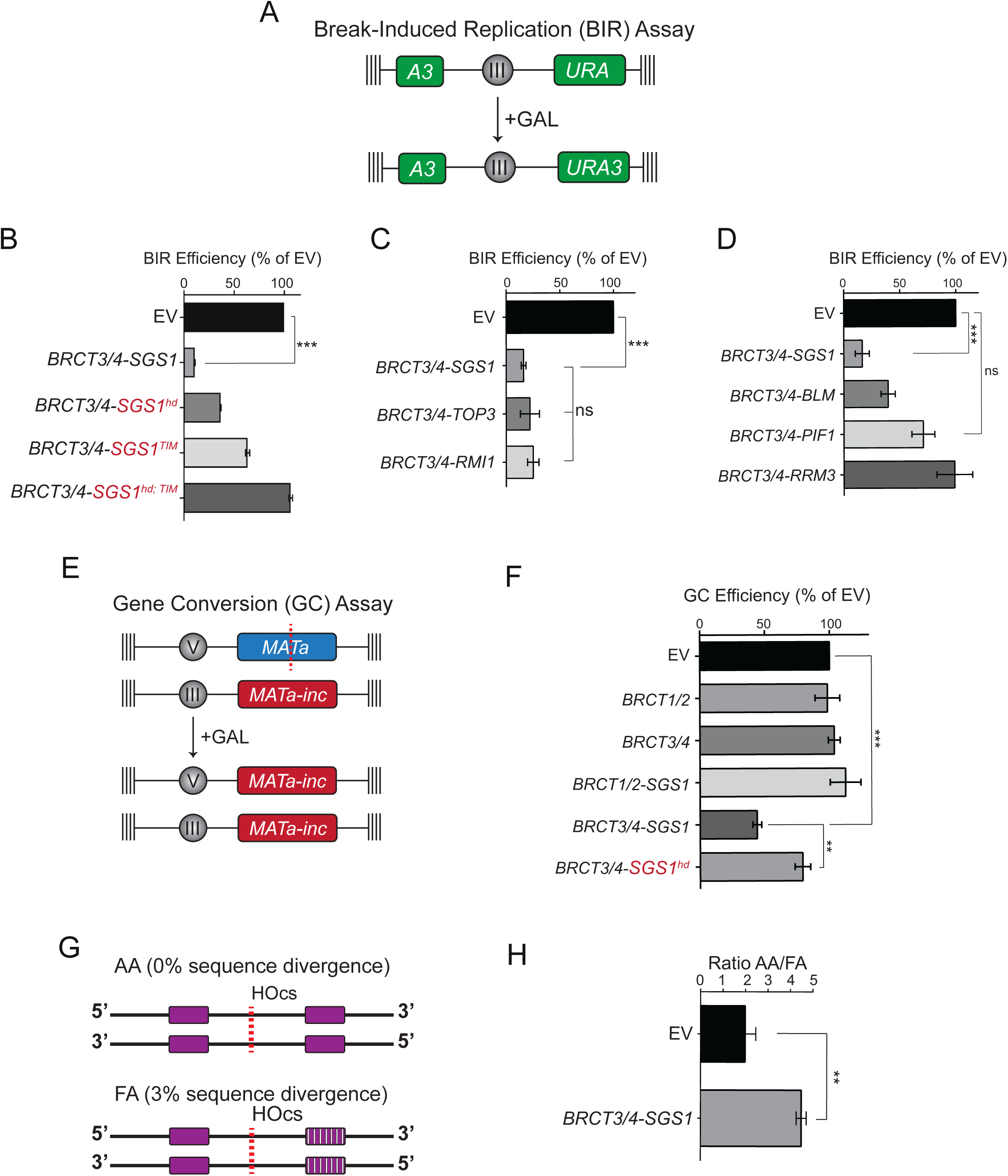
The B3/4-Sgs1 Fusion Protein Impairs Homologous Recombination. A. Diagram of the break-induced replication (BIR) assay used in this study. B. Measurement of BIR efficiency in cells expressing empty vector or BRCT3/4 of Dpb11 fused to wild-type or mutant versions of Sgs1. Mutants include a helicase deficient allele of Sgs1 (*SGS1*^*hd*^) or an allele lacking the critical amino acid residues responsible for binding to Top3 (*SGS1*^*TIM*^). Error bars represent SEM of at least 3 replicate experiments. C. Measurement of BIR efficiency in cells expressing empty vector or BRCT3/4 of Dpb11 fused to either SGS1, TOP3, or RMI1. Error bars represent SEM of at least 3 replicate experiments. D. Measurement of BIR efficiency in cells expressing empty vector or BRCT3/4 of Dpb11 fused to either SGS1, hBLM, PIF1, or RRM3. Error bars represent SEM of at least 3 replicate experiments. E. Diagram of the gene conversion (GC) assay used in this study. F. Measurement of gene conversion efficiency in cells expressing empty vector, BRCT1/2 of Dpb11, BRCT3/4 of Dpb11, or BRCT3/4 of Dpb11 fused to the helicase defective allele of SGS1 (*SGS1*^*hd*^). Error bars represent SEM of at least 3 replicate experiments. G. Diagram of the single-strand annealing (SSA) assay used in this study. H. Measurement of single-strand annealing in cells expressing empty vector or BRCT3/4-SGS1. Error bars represent SEM of at least 3 replicate experiments.

## DISCUSSION

Accumulated evidence over the past 30 years has identified Mec1 as a central player in the preservation of genome stability, particularly through the regulation of distinct steps of HR such as DNA end resection (Lanz et al., 2019). However, the precise mechanisms and targets regulated by Mec1 signaling for ensuring proper HR control remain incompletely understood, representing a major knowledge gap in our understanding of genome maintenance mechanisms. In this work, we sought to identify novel connections between Mec1/ATR signaling and the homologous recombination machinery. By using *rad9Δ* cells, we aimed at analyzing a condition in which DNA lesions become hyper-resected and HR is dysregulated at a key early step. We reasoned that the scenario of extensive ssDNA accumulation should increase the representation of recombination intermediates and Mec1-dependent signaling events important for HR control that are often too transient in wild type cells. Consistent with the extensive processing of DNA ends in *rad9Δ* cells, our phosphoproteomic results revealed that in these cells Mec1/ATR preferentially targets proteins involved in ssDNA-associated transactions, including Rfa2, Uls1 and Sgs1. Therefore, the use of *rad9Δ* cells and quantitative MS analysis of phosphosignaling events allowed us to capture a mode of Mec1/ATR signaling that is under-represented in wild-type cells. We propose that this specific mode of Mec1/ATR signaling is important to control HR-mediated repair events. Taken together, the identification of a distinct mode of Mec1 signaling triggered by hyper-resection highlights the multi-faceted action of this kinase in the coordination of checkpoint signaling and HR-mediated DNA repair (see model in Fig. 8).

**Figure 8.**
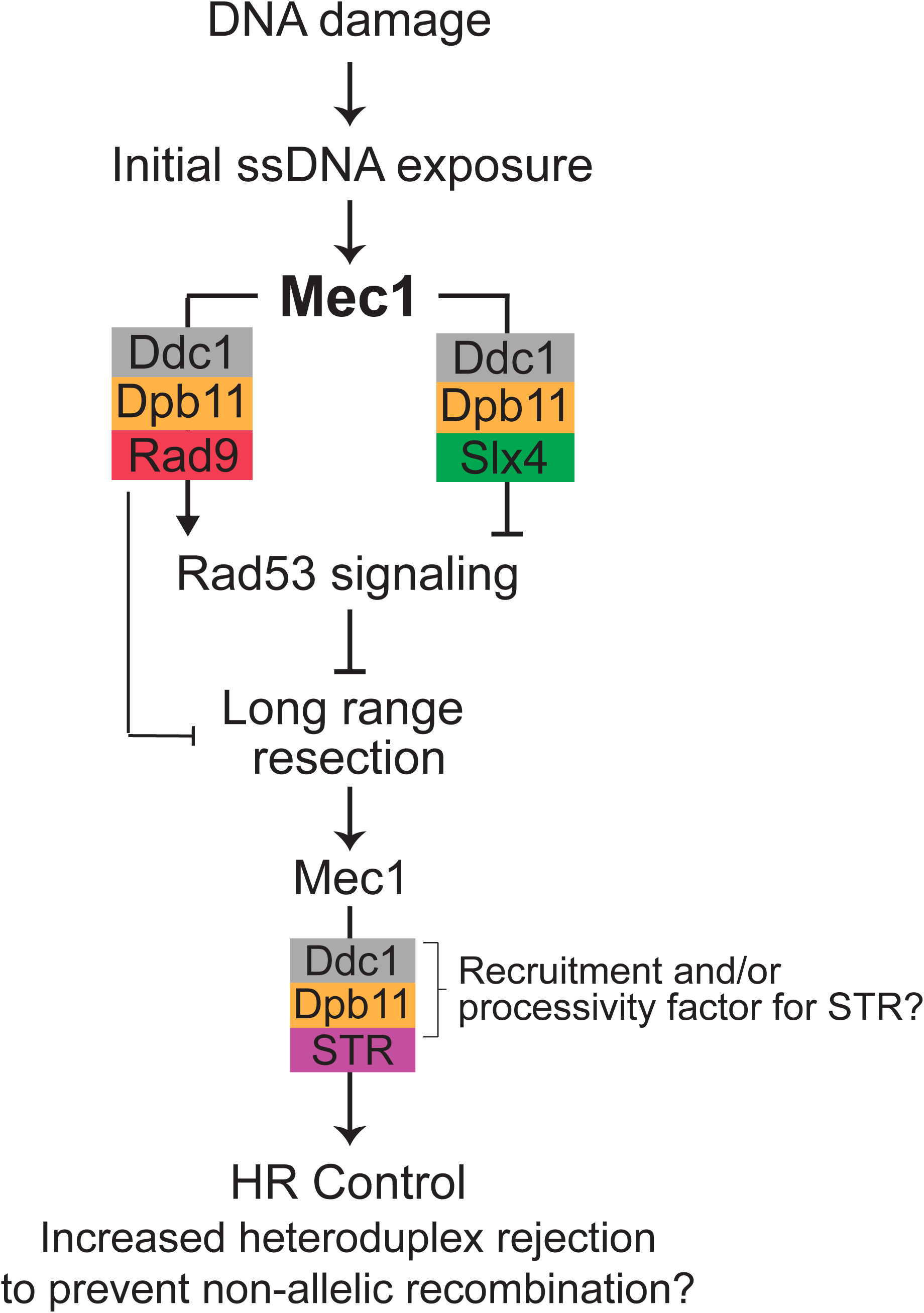
Model for Distinct Modes of Mec1 Signaling in the Control of Checkpoint Signaling and Homologous Recombination. Mec1 is recruited to RPA-ssDNA following DNA damage to promote Rad53-mediated checkpoint signaling that prevents immediate long-range resection of DNA ends. This anti-resection function of Mec1 is important to protect DNA ends and ensure that subsequent resection occurs in a restrained and controllable manner. Since the Dpb11-Rad9 interaction is not dependent on Mec1 (but mostly dependent on CDK phosphorylation), the Dpb11-Rad9 complex is rapidly stabilized at DNA lesions once initial Mec1 signaling and Ddc1 phosphorylation occurs. As Mec1 signaling builds up, it hyper-phosphorylates the Slx4 scaffold, which becomes a strong interactor of Dpb11 and displaces Rad9 to counteract checkpoint signaling and Rad9-mediated resection block. Mec1 signaling therefore switches to an anti-checkpoint and pro-resection mode. Once long-resection occurs, extensive ssDNA accumulates, leading to increased opportunities for strand invasion, but also increased opportunities for non-allelic recombination (*rad9Δ* cells have increased non-allelic recombination, (Fasullo, Bennett, Ahching, & Koudelik, 1998; Nielsen, Bentsen, Andersen, Gasser, & Bjergbaek, 2013)). In this context, we propose that a new mode of Mec1 signaling triggered by extensive resection stabilizes the STR complex at lesions via interaction with Dpb11 for proper regulation of HR. It is tempting to speculate that the Dpb11-9-1-1 complex acts as a “processivity factor” for the helicase function of Sgs1 to processively displace, and reject, heteroduplexes.

In addition to identifying a novel mode of Mec1/ATR signaling triggered by hyper-resection, our work finds that such mode of signaling converges toward the assembly of a novel protein complex between the STR complex and the Dpb11 scaffold. In *rad9Δ* cells, Mec1/ATR heavily phosphorylates the STR subunit Sgs1. While we find Mec1 signaling to be required for assembly of the STR-Dpb11, several details of how the multi-phosphorylation pattern in Sgs1 contributes to the interaction with Dpb11 remain incompletely understood. We were unable to identify a specific phosphorylation site, or a set of phosphorylation events, required for mediating the interaction. The interaction may involve additional kinases downstream of Mec1, since mutation of all S/T-Q sites on Sgs1 did not ablate Sgs1 binding to Dpb11. Alternatively, phosphorylation of any S/T-Q residue on Sgs1, Top3, or Rmi1 may be sufficient to induce the formation of STR-Dpb11 complexes. It also remains possible that phosphorylation of Ddc1 threonine 602 is the key Mec1-dependent phosphorylation important for mediating the Dpb11-Sgs1 interaction. In this scenario, the extensive exposure of ssDNA may promote the selective recruitment of STR complex close to Dpb11-9-1-1 complex, which is then recognized by BRCT1/2 of Dpb11 in a phosphorylation-independent manner. We don’t favor this model, since BRCT1/2 of Dpb11 is well known to interact with phosphorylated epitopes (Bork et al., 1997; Cussiol et al., 2015; Pfander & Diffley, 2011; Yu, Chini, He, Mer, & Chen, 2003). Of importance, we don’t exclude the possibility that most of Mec1-dependent phosphorylation events in Sgs1 are not involved in mediating the interaction with Dpb11, and that those may regulate additional aspects of Sgs1 function, such as conformational changes that alter its activity or ability to interact with other proteins.

Sgs1 is a well-established regulator of both early and later stages of homology-directed repair (Ira et al., 2003; Mankouri, Ashton, & Hickson, 2011; Sugawara et al., 2004; Zhu, Chung, Shim, Lee, & Ira, 2008). Sgs1 has well defined roles in monitoring and disassembling recombination intermediates, including recombination-driven heteroduplexes (Cejka, Plank, Dombrowski, & Kowalczykowski, 2012). Therefore, our findings suggested the model that upon hyper-resection of DNA ends, Mec1/ATR signaling converges to Sgs1 (and STR complex) as part of a quality control mechanism to prevent the aberrant exposure of ssDNA from triggering erroneous HR events. Although alternative models remain plausible, the finding that the BRCT3/4-Sgs1 fusion protein impairs HR-mediated repair supports the quality control model. Of importance, such fusion did not alter Rad53 signaling, congruent with the idea that the chimera does not interfere with checkpoint signaling control. Since Rad53 signaling is often tightly correlated with the extent of resection, this finding suggests that resection is not affected by expression of the B3/4-Sgs1 chimera. This is not surprising, since Sgs1 requires Dna2 in its function of promoting long range resection (Zhu et al., 2008). As such, the results are consistent the model of the chimera impairing heteroduplex stability, and due to its strong dominant effect, preventing recombination events at large. Interestingly, experiments using the B3/4-BLM^Human^ chimera suggest a conserved role for the STR-Dpb11 interaction in regulating recombination. Indeed, BLM and TOPBP1 were reported to interact in human cells (Blackford et al., 2015; Sun et al., 2017), although it is unclear whether ATR plays any role in promoting that interaction, and if the BLM-TOPBP1 interaction affects HR.

In addition to STR phosphorylation, we also detected that lack of *RAD9* induces the phosphorylation of Uls1, a ubiquitin ligase and DNA translocase also reported to control HR. While less is understood about Uls1 compared to Sgs1, genetic studies have linked Uls1 to Sgs1 through Uls1’s translocase function, which may channel recombination intermediates into Sgs1-dependent repair mechanisms (Cal-Bkowska, Litwin, Bocer, Wysocki, & Dziadkowiec, 2011). Future work focused on the role of Mec1-phosphorylation of Uls1 may illuminate further connections between Uls1, Sgs1 and Mec1, and potentially reveal novel mechanisms regulating recombination in response to hyper-resection. Interestingly, Uls1 did appear in our Dpb11 IP-MS experiments as a Dpb11 interactor in *rad9*Δ cells, but the number of peptides identified was too low to pass our threshold for calling Dpb11 interactors.

Taken together, our data uncovered a distinct mode of Mec1 signaling in response to hyper-resection that targets key factors involved in the regulation of HR-mediated repair. The discovery of the Dpb11-STR complex and its mode of interaction and engagement with the 9-1-1 complex adds an important additional role for Dpb11 in controlling the DNA damage response. Notably, the assembly of the complex and requirements for interaction follow largely a similar logic to what was previously reported for the Dpb11-Rad9 and Dpb11-Slx4-Rtt107 complexes (Cussiol et al., 2015; Gritenaite et al., 2014; Ohouo et al., 2010; Pfander & Diffley, 2011). Future work should focus on elucidating how the formation of the three independent Ddc1-dependent Dpb11 complexes (Slx4-Dpb11, Rad9-Dpb11, and Sgs1-Dpb11) are spatiotemporally regulated. For example, it would be interesting to test whether Dpb11 interactors compete directly or instead bind discrete 9-1-1 complexes. Biochemical data suggests that 9-1-1 clamps, once loaded, can slide along plasmids DNA substrates, on both ssDNA and dsDNA (Majka & Burgers, 2003), raising the possibility that many 9-1-1 clamps could be loaded onto DNA per lesion, with each clamp harboring a Dpb11 molecule and its associated interacting partner, dependent perhaps on the sequence or structural context of the DNA. Future work may also reveal novel Mec1-dependent interactors of Dpb11, expanding on the common logic for how Mec1 coordinates the DNA damage response and cementing Dpb11 as a critical scaffolding protein which integrates Mec1 signaling inputs into the formation of concerted repair protein complex outputs.

## MATERIALS AND METHODS

### Yeast Strains

A complete list of yeast strains used in this study can be found in Supplemental Table S5. The background for all yeast used in this study is S288C. Whole ORF deletions were performed using the established PCR-based strategy to amplify resistance cassettes with flanking sequence homologous to a target gene. All whole ORF deletions were verified by PCR. Primer sequences for gene deletions are available in Supplemental Table S5. Yeast were grown at 30°C in either rich or synthetic dropout media depending on the experiment. Plasmids in this study are listed in Supplemental Table S5 and are available upon request.

### Co-Immunoprecipitation (Co-IP)

Yeast cell lysates were prepared for immunoprecipitation by bead beating for 3 cycles of 10 minutes with 1-minute rest between cycles at 4°C in lysis buffer (150mM NaCl, 50mM Tris pH 7.5, 5mM EDTA, 0.2% Tergitol type NP40) supplemented with complete EDTA-free protease inhibitor cocktail (Roche), 5 mM sodium fluoride and 10 mM β-glycerophosphate. Following normalization by Bradford assay, ∼5mg of lysate per IP was incubated with antibody-conjugated agarose resin for 3 hours at 4°C. Resin was washed 4x in lysis buffer. Elution was performed either with FLAG peptide or elution buffer (1% SDS, 100mM Tris pH 8.0).

### Immunoprecipitation-Mass Spectrometry (IP-MS)

For IP-MS experiments, control yeast or yeast expressing tagged bait proteins were grown in “heavy” or “light” SILAC media [complete synthetic medium -Arg-Lys supplemented with either isotopically heavy (containing C^13^ and N ^15^) or normal (containing C^12^ and N ^14^) lysine and arginine] to mid-log phase and treated as described in the figure legend. Cells were pelleted at 1000rcf and washed with TE buffer containing 1mM PMSF. Pellets from “light” and “heavy” samples were lysed and processed separately as described for Co-IP above. Proteins bound to antibody-conjugated agarose resin were eluted with 1% SDS, 100mM Tris pH 8.0, and then “light” and “heavy” eluates were mixed, reduced with 10mM DTT and alkylated with 25mM iodoacetamide followed by precipitation on ice for 1hr in PPT solution (50% acetone, 49.9% ethanol, 0.1% acetic acid). Pellets were washed once with PPT then resuspended in urea/tris solution (8M urea, 50mM Tris pH 8.0). Urea-solubilized pellet was then diluted to 2M urea using NaCl/Tris solution (150mM NaCl, 50mM Tris pH 8.0) and digested overnight at 37°C with 10ug of trypsin GOLD (Promega). The following day, samples were desalted using a 50mg Waters Sep-Pak column. Eluted peptides were dried and resuspended in 0.1% TFA and subjected for LC-MS/MS analysis on a Thermo-Fisher Q-Exactive HF mass spectrometer as recently described (Lanz et al., 2018).

### Phosphoproteomics

For phosphoproteomic experiments, 200mL of yeast grown in “heavy” or “light” SILAC media [complete synthetic medium -Arg-Lys supplemented with either isotopically heavy (containing C^13^ and N ^15^) or normal (containing C^12^ and N ^14^) lysine and arginine] to mid-log phase and treated as described in the figure legend. Cells were pelleted at 1000rcf and washed with TE buffer containing 1mM PMSF. Pellets were lysed by bead beating with 0.5mm glass beads for 3 cycles of 10 minutes with 1-minute rest time between cycles at 4°C in lysis buffer (150mM NaCl, 50mM Tris pH 7.5, 5mM EDTA, 0.2% Tergitol type NP40) supplemented with complete EDTA-free protease inhibitor cocktail (Roche), 5 mM sodium fluoride and 10 mM β-glycerophosphate. Seven mg of each light and heavy labeled protein lysate was denatured and reduced with 1% SDS and 5mM DTT at 65°C, then alkylated with 25mM iodoacetamide. Light and heavy protein lysates were mixed and precipitated with a cold solution of 50% acetone, 49.9% ethanol, 0.1% acetic acid. Protein pellet was resuspended with 2M urea and subsequently digested with TPCK-treated trypsin overnight at 37°C. Phosphoenrichment was performed using a Thermo-Fisher Fe-NTA phosphopeptide enrichment kit (cat# A32992) according to the manufacturer’s protocol. Purified phosphopeptides were then fractionated using HILIC chromatography and subjected to LC-MS/MS on a Thermo-Fisher Q-Exactive HF mass spectrometer as recently described (Lanz et al., 2018).

### Mass Spectrometry Data Analysis

For IP/MS experiments, raw MS/MS spectra were searched using the SORCERER (Sage N Research, Inc.) system running the SEQUEST software over a composite yeast protein database, consisting of both the normal yeast protein sequences downloaded from the Saccharomyces Genome Database (SGD) and their reversed protein sequences as a decoy to estimate the false discovery rate (FDR) in the search results. Searching parameters included a semi-tryptic requirement, a mass accuracy of 15 ppm for the precursor ions, differential modification of 8.0142 daltons for lysine, 10.00827 daltons for arginine, and a static mass modification of 57.021465 daltons for alkylated cysteine residues. XPRESS software, part of the Trans-Proteomic Pipeline (Seattle Proteome Center), was used to quantify all the identified peptides. Proteins with fewer than 4 PSMs identified were excluded. Statistical analyses were performed using a Wilcoxon rank-sum test.

For phosphoproteomic experiments, raw MS/MS spectra were searched using the COMET engine (part of the Trans-Proteomic Pipeline; Seattle Proteome Center) over a composite yeast protein database, consisting of both the normal yeast protein sequences downloaded from the Saccharomyces Genome Database (SGD) and their reversed protein sequences as a decoy to estimate the false discovery rate (FDR) in the search results. Searching parameters included a semi-tryptic requirement, a mass accuracy of 15 ppm for the precursor ions, differential modification of 8.0142 daltons for lysine, 10.00827 daltons for arginine, 79.966331 daltons for phosphorylation of serine, threonine and tyrosine (for phosphoproteomic experiments) and a static mass modification of 57.021465 daltons for alkylated cysteine residues. Phosphorylation site localization probabilities were determined using PTMProphet and quantitation of identified phosphopeptides was performed using XPRESS (both tools part of the Trans-Proteomic Pipeline; Seattle Proteome Center). Phosphoproteomic data shown in Figures 1 and 2 represent the combined independent results of a “forward” (condition 1 = light / condition 2 = heavy) and a “reverse” (condition 1 = heavy / condition 2 = light) SILAC experiment. Using this experimental design, phosphorylation events that were not consistently identified in two independent, separately SILAC labelled yeast cultures were filtered out. All mass spectrometric data presented in this study are available through PRIDE.

### Western Blots

Yeast cell lysates were prepared for western blotting by bead beating for 15 minutes at 4°C in lysis buffer (150mM NaCl, 50mM Tris pH 7.5, 5mM EDTA, 0.2% Tergitol type NP40) supplemented with complete EDTA-free protease inhibitor cocktail (Roche), 5 mM sodium fluoride and 10 mM β-glycerophosphate. Following normalization by Bradford assay, lysates were boiled in Laemmli buffer and electrophoresed on a 9% SDS-PAGE gel. Proteins were transferred wet onto a PVDF membrane and incubated with antibody. Signal detection was performed using HRP-coupled secondary antibodies in all cases, imaged either with a BioRad ChemiDoc or X-ray film.

### Genetic Assays to Measure Recombination

BIR assay was performed according to (Anand et al., 2014). GC assay was performed as described in (Ira et al., 2003). SSA assay was performed according to (Sugawara et al., 2004).

### Spot Assays

For dilution assays, 5mL of yeast culture was grown to saturation at 30°C. Then, 1 OD600 equivalent of the saturated culture was 10-fold serially diluted in a 96-well plate in water and spotted onto agar plates using a bolt pinner.

### Data Availability

The mass spectrometry data from this publication have been deposited to the PRIDE database (https://www.ebi.ac.uk/pride/archive/) and assigned the identifiers PXD017286 and PXD017289.

## ACKNOWLEDGEMENTS

We thank Beatriz S. Almeida for technical support; we thank Jim Haber, Gregorz Ira, and Eric Alani for graciously providing yeast strains YRA213, TGI354, and EAY1141/54, respectively; we thank Xiaolan Zhao for careful reading of the manuscript and comments; and we thank Diego Dibitetto and other members of the Smolka lab for valuable discussions related to this work. This work is supported by a grant from the National Institute of Health (R01-GM097272) to M.B.S.

## AUTHOR CONTRIBUTIONS

EJS and MBS conceptualized the project, designed experiments, interpreted data, and wrote the paper. EJS, SCV, and WJC performed experiments. VMC and MBS designed the phosphoproteomics data analysis pipeline. All authors contributed to analyzing data and revising the manuscript.

